# Biofilm-associated toxin and extracellular protease cooperatively suppress competitors in *Bacillus subtilis* biofilms

**DOI:** 10.1101/663328

**Authors:** Kazuo Kobayashi, Yukako Ikemoto

**Affiliations:** Division of Biological Science, Nara Institute of Science & Technology

**Keywords:** antibiotics, toxin/anti-toxin, protease, biofilm, biofilm matrix

## Abstract

In nature, most bacteria live in biofilms where they compete with their siblings and other species for space and nutrients. Some bacteria produce antibiotics in biofilms; however, since the diffusion of antibiotics is generally hindered in biofilms by extracellular polymeric substances, i.e., the biofilm matrix, their function remains unclear. The *Bacillus subtilis yitPOM* operon is a paralog of the *sdpABC* operon, which produces the secreted peptide toxin SDP. Unlike *sdpABC*, *yitPOM* is induced in biofilms by the DegS-DegU two-component regulatory system. High *yitPOM* expression leads to the production of a secreted toxin called YIT. Expression of *yitQ*, which lies upstream of *yitPOM*, confers resistance to the YIT toxin, suggesting that YitQ is an anti-toxin protein for the YIT toxin. The alternative sigma factor SigW also contributes to YIT toxin resistance. In a mutant lacking *yitQ* and *sigW*, the YIT toxin specifically inhibits biofilm formation, and the neutral protease NprB is required for this inhibition. The requirement for NprB is eliminated by Δ*eps* and Δ*bslA* mutations, either of which impairs production of biofilm matrix polymers. Overexpression of biofilm matrix polymers prevents the action of the SDP toxin but not the YIT toxin. These results indicate that, unlike the SDP toxin and conventional antibiotics, the YIT toxin can pass through layers of biofilm matrix polymers to attack cells within biofilms with assistance from NprB. When the wild-type strain and the YIT-sensitive mutant were grown together on a solid medium, the wild-type strain formed biofilms that excluded the YIT-sensitive mutant. This observation suggests that the YIT toxin protects *B. subtilis* biofilms against competitors. We propose that some bacteria have evolved specialized antibiotics that can function within biofilms.

**Author Summary:** Biofilms are multicellular aggregates of bacteria that are formed on various living and non-living surfaces. Biofilms often cause serious problems, including food contamination and infectious diseases. Since bacteria in biofilms exhibit increased tolerance or resistance to antimicrobials, new agents and treatments for combating biofilm-related problems are required. In this study, we demonstrated that *B. subtilis* produces a secreted peptide antibiotic called the YIT toxin and its resistant protein in biofilms. A mutant lacking the resistance gene was defective in biofilm formation. This effect resulted from the ability of the YIT toxin to pass through the biofilm defense barrier and to attack biofilm cells. Thus, unlike conventional antibiotics, the YIT toxin can penetrate biofilms and suppress the growth of YIT toxin-sensitive cells within biofilms. Some bacteria produce antibiotics in biofilms, some of which can alter the bacterial composition in the biofilms. Taking these observations into consideration, our findings suggest that some bacteria produce special antibiotics that are effective against bacteria in biofilms, and these antibiotics might serve as anti-biofilm agents.

## Introduction

In the environment, bacteria compete for space and nutrients [1]. Antibiotics are thought to play a critical role in this competition, and antibiotic-producing bacteria are indeed common in various environments [2–4]. In the environment, most bacteria are found in sessile multicellular bacterial communities known as biofilms, in which the cells benefit from increased antibiotic tolerance or resistance [5, 6]. Though alternative environmental roles of antibiotics have been proposed [7], this paradox has not been explained in detail to date.

In biofilms, cells adhere to each other and to a surface via a mixture of extracellular polymeric substances called the biofilm matrix, which consists of exopolysaccharides, proteins, nucleic acids, and/or lipids [8, 9]. When encased in the biofilm matrix, cells exhibit increased tolerance or resistance to environmental stresses, antibiotics, host defense systems, and predation [8, 9]. Thus, biofilm formation enables bacteria to remain in a favorable niche and to claim territory; however, biofilms are not a utopia for bacteria. The properties of biofilms, including high cell density, decreased internal fluidity, and, in many cases, the presence of multiple species, lead to conditions of harsh competition, especially when nutrients are scarce. Many bacteria secrete biofilm formation-inhibiting molecules, such as biosurfactants, polysaccharides, and molecules that interfere with bacterial quorum sensing, and these secreted molecules help to exclude unfavorable competitors from biofilms [10]. Antibiotics might also play an important role in competition within biofilms. However, since the properties of biofilms, including the protection of member cells by the biofilm matrix, the increased antibiotic tolerance, and the decreased internal fluidity, reduce the efficacy of antibiotics against biofilm cells [11–16], little attention has been paid to the functions of antibiotics in competition within biofilms. However, some biofilms do indeed produce antibiotics, and several of these antibiotics can alter the bacterial composition of the biofilm [17–23]. These observations suggest that the functions of antibiotics produced in biofilms remain to be investigated. An understanding of how bacteria use antibiotics in biofilms will not only provide insight into bacterial survival strategies within biofilms, it will also lead to the discovery of tactics for combating biofilm-related problems, such as food and beverage safety issues, industrial contamination, and infectious diseases.

The Gram-positive soil bacterium *Bacillus subtilis* is a model organism for biofilm formation. *B. subtilis* forms robust biofilms under laboratory conditions, for example, pellicles on the surface of liquid media under static culture conditions or wrinkled colonies on solid media [24]. *B. subtilis* biofilms are maintained by a biofilm matrix that mainly consists of exopolysaccharides, TasA amyloid fibers, and BslA hydrophobins, which are produced by proteins encoded by the *epsABCDEFGHIJKLMNO* operon, the *tapA-sipW-tasA* operon, and *bslA*, respectively [24–29]. These genes are directly or indirectly repressed by the transcriptional repressors AbrB and SinR [30–33]. Phosphorylation of the response regulator Spo0A induces mechanisms that antagonize these repressors, leading to the expression of the biofilm matrix synthesis genes [34, 35].

*B. subtilis* produces a wide array of antibiotics. Many of these antibiotics are non-ribosomally synthesized peptide compounds, such as surfactin, bacillaene, fengycin, iturin, and bacilysin, which are thought to be important in nature for competition with other organisms, including fungi [4, 36]. Furthermore, *B. subtilis* produces ribosomally synthesized peptide antibiotics, such as bacteriocins and other protein-derived toxins, which are generally effective against other bacteria that are genetically similar and present in similar ecological niches [4, 37–39]. One of these protein-derived toxins is the cannibalism toxin SDP [40], whose function is involved in biofilm formation. The SDP toxin is derived from the internal sequence of SdpC, and it is encoded by the *sdpABC* operon. SdpC is a 203 amino acid protein that contains a typical N-terminal secretion signal and a C-terminal hydrophobic domain. After secretion and cleavage of the signal sequence, SdpC is further processed into the 42 amino acid peptide known as the SDP toxin, which corresponds to the C-terminal hydrophobic domain (C141 to S182) [39–41]. SdpA and SdpB are required for the processing of SdpC to SDP, and this processing is essential for the activity of the SDP toxin [41]. The SDP toxin penetrates bacterial membranes, where it then induces cell lysis by collapsing the proton motive force [42]. Downstream of *sdpABC* is the *sdpRI* operon, which encodes its own transcriptional repressor and an anti-toxin protein to the SDP toxin [43]. SdpI is an integral membrane protein that protects cells probably by binding to the SDP toxin. Transcription of the *sdpABC* and *sdpRI* operons is directly or indirectly activated by phosphorylated Spo0A [40, 43]. Spo0A is a master regulator of stationary phase development that is phosphorylated after the onset of stationary phase [44]. However, as the phosphorylation of Spo0A is subject to a bistable regulatory mechanism, a subset of *B. subtilis* cells produce the SDP toxin and the SdpI anti-toxin protein [40, 43]. Consequently, the secreted SDP toxin lyses and kills a fraction of the sibling cells that do not produce the SdpI anti-toxin protein. Since phosphorylated Spo0A also induces biofilm formation in parallel, cells that produce the SDP toxin efficiently develop biofilms by using nutrients released from their lysed siblings [45]. Moreover, the SDP toxin is effective not only against *B. subtilis*, but also against many Firmicutes bacteria [39, 41, 46]. Thus, the SDP toxin likely plays an important role in the early phase of biofilm formation by eliminating unnecessary types of cells and closely related competitors in the environment.

The undomesticated *B. subtilis* strain NCIB3610 encodes an *sdpABC* paralog known as *yitPOM*. YitP and YitO exhibit approximately 50% similarity to the entire SdpA and SdpB sequences, respectively (S1A Fig). Like SdpC, YitM has an N-terminal secretion signal; however, the sequence similarity between YitM and SdpC is limited to the N-terminal three quarters of the sequence, which does not include the entire sequence corresponding to the SDP toxin (S1B Fig). Although there is no sequence similarity, like SdpC, the YitM C-terminal region contains a hydrophobic domain (S1C Fig). These observations suggest the possibility that the C-terminal hydrophobic domain of YitM might be processed to a secreted toxin via a YitP and YitO-dependent mechanism. If this is the case, then *yitPOM* encodes a toxin whose sequence differs from that of the SDP toxin. While transcription of *sdpABC* is activated by Spo0A [40], *yitPOM* was previously identified as a member of the group of genes regulated by the DegS-DegU two-component regulatory system [47]. Since DegS-DegU regulates biofilm formation [47, 48], we were interested in determining the functions of the *yitPOM*-encoded toxin, particularly in biofilms.

In this paper, we demonstrate that *yitPOM* encodes a biofilm-specific secreted toxin. Unlike conventional antibiotics, this toxin could attack cells within biofilms by passing through the layers of the biofilm matrix polymers with the assistance of an extracellular protease. Our results suggest that some bacteria have evolved specialized antibiotics to suppress the growth of competitors in biofilms.

## Results

### *yitPOM* encodes a toxin

We investigated the function of the *yitPOM* operon using *B. subtilis* strain NCIB3610 (hereafter referred to as the wild-type strain or 3610) [24]. To determine whether *yitPOM* encodes a toxin, we examined the effect of *yitPOM* overexpression on growth. We constructed the strain P*_spac_*_-hy_-*yitPOM*, which ectopically expresses *yitPOM* from the strong isopropyl β-D-thiogalactopyranoside (IPTG)-inducible, LacI-repressible *spac*-hy promoter [49] in the *amyE* locus on the chromosome. The wild-type and P*_spac_*_-hy_-*yitPOM* strains were grown with vigorous shaking in 2× Schaeffer’s sporulation medium plus glucose (2×SG) [50] supplemented with 1 mM IPTG, and the optical density at 600 nm (OD_600_) was measured over time. These strains showed no difference in growth from exponential phase to stationary phase (Fig 1A). Toxin-producing bacteria normally express cognate anti-toxin proteins against their toxins, and the effects of the toxins do not appear unless the anti-toxin genes are deleted [2, 37, 38, 40]. Since the toxin and anti-toxin genes are frequently located close to each other in the genome [2, 37, 38, 40], genome comparison is a powerful tool to identify toxin/anti-toxin gene sets. To identify candidates for an anti-toxin gene against the putative toxin encoded by *yitPOM*, we compared the genetic organization of the 3610 and *B. subtilis* var. *natto* BEST195 strains, the latter of which lacks *yitPOM*. This comparison revealed that *yitPOM* appears to be inserted between *yitR* and *yitL* in the 3610 genome, along with *yizB* and *yitQ*, which encode a transcriptional regulator and a membrane protein, respectively (Fig 1C). Since the SdpI anti-toxin protein is a membrane protein, YitQ was a candidate for an anti-toxin protein to the putative toxin, although YitQ has no similarity to SdpI. Based on the DNA sequence, *yizB* and *yitQ* are predicted to form an operon with a downstream gene, *yitR*, which also encodes a membrane protein. Althogh *yitR* is present in BEST195, we kept it as a second candidate for the anti-toxin protein. We were concerned that deleting these candidate anti-toxin genes might cause severe growth defects by releasing the activity of the toxin encoded by the genomic *yitPOM* operon. Therefore, we constructed a Δ*yitR-yitM* deletion strain that lacks the entire region from *yitR* to *yitM*, which contains the candidate anti-toxin genes *yitR* and *yitQ*, the unknown repressor gene *yizB*, and the putative toxin-encoding *yitPOM* operon (Fig 1C). Furthermore, since the expression of the secondary resistance mechanism against the SDP toxin is induced by the alternative sigma factor SigW (σ^W^) [51], we also constructed a *sigW* deletion strain. Subsequently, either the Δ*yitR-yitM* and Δ*sigW* mutation alone or both mutations together were introduced into the P*_spac_*_-hy_-*yitPOM* strain. We compared the growth of these strains in 2×SG medium supplemented with 1 mM IPTG. While the P*_spac_*_-hy_-*yitPOM* Δ*yitR-yitM* and P*_spac_*_-hy_-*yitPOM* Δ*sigW* mutants grew normally, the P*_spac_*_-hy_-*yitPOM* Δ*yitR-yitM* Δ*sigW* mutant showed mild cell lysis 3 h after the end of exponential phase (Fig 1A). We confirmed that *yitPOM* expression caused this cell lysis, as cell lysis was only observed when *yitPOM* expression was induced with IPTG (Fig 1B). These results indicate that *yitPOM* expression leads to the production of a toxin that causes cell lysis in a mutant strain lacking the putative anti-toxin genes and *sigW*.

**Fig 1.**
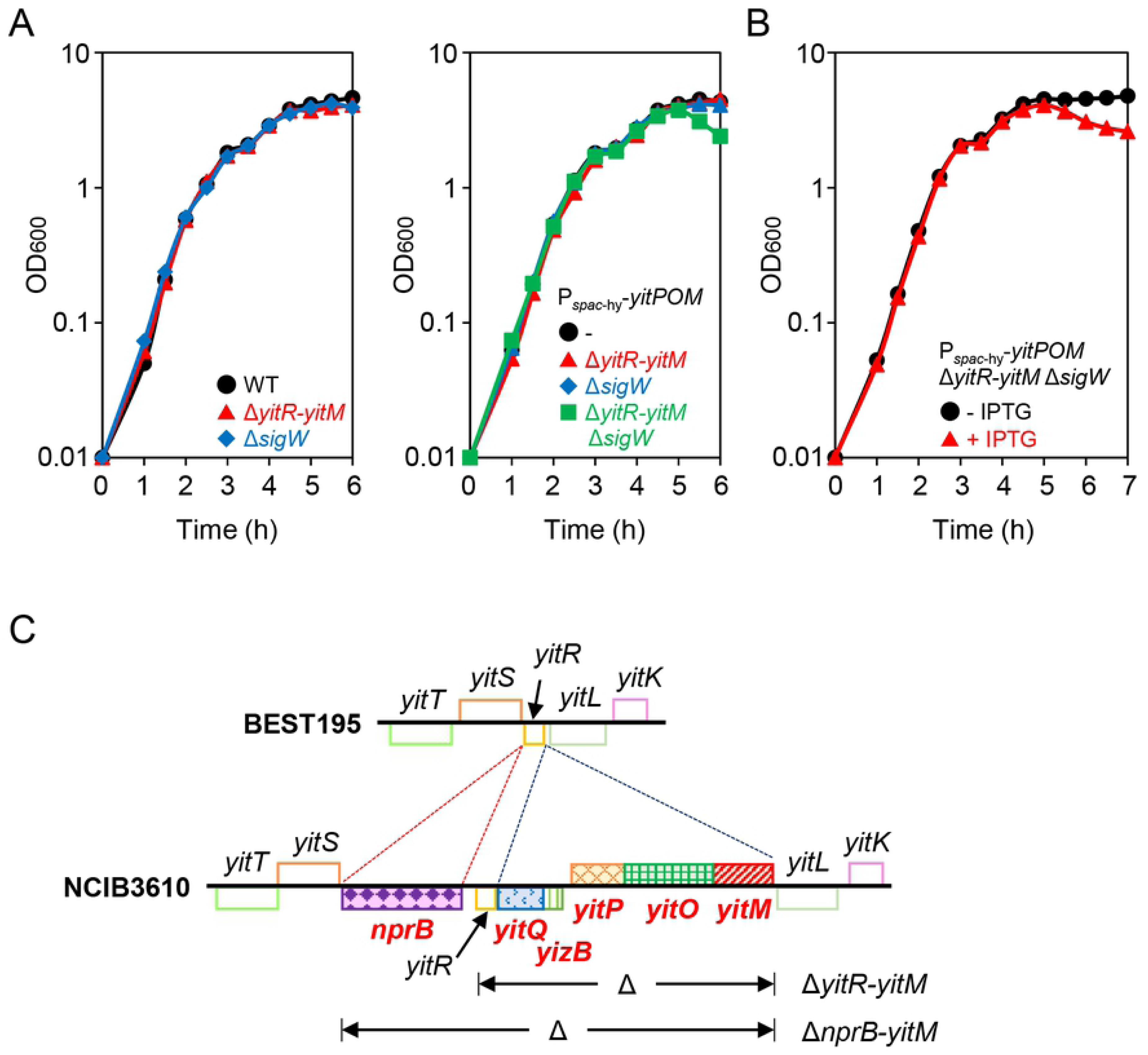
*yitPOM* encodes a toxin. (A) Effect of *yitPOM* induction on cell growth. *B. subtilis* strains were grown at 37°C in 2×SG supplemented with 1 mM IPTG with vigorous shaking. (B) Induction of *yitPOM* caused mild cell lysis. The P*_spac_*_-hy_-*yitPOM* Δ*yitR-yitM* Δ*sigW* strain was grown in 2×SG supplemented with or without 1 mM IPTG. (C) Comparison of the genetic organization in NCIB3610 and BEST195. Homologous genes are shown by boxes of the same color. Genes only present in NCIB3610 are shown in red bold. The deleted regions in the Δ*yitR-yitM* and Δ*nprB-yitM* mutants are shown below the gene map of NCIB3610.

To further confirm that *yitPOM* encodes a toxin, we employed a spot-on-lawn assay. We performed this assay in the Δ*sdpABC-sdpIR* (hereafter referred to as Δ*sdpA-sdpR*) Δ*yitR-yitM* mutant background to eliminate the effects of the endogenous *sdpABC* and *yitPOM* operons. Since *spo0A* null mutants are sensitive to multiple antibiotics and stresses, including the SDP toxin [40, 52, 53, 54], we used the Δ*sdpA-sdpR* Δ*yitR-yitM* Δ*spo0A* mutant as an antibiotic-sensitive indicator strain. When spotted on a lawn of this indicator strain, the strain expressing YitPOM (P*_spac_*_-hy_-*yitPOM* Δ*sdpA-sdpR* Δ*yitR-yitM*) formed growth inhibition zones (halos) around its colonies (Fig 2). By contrast, a strain that does not express YitPOM (Δ*sdpA-sdpR* Δ*yitR-yitM*) formed no obvious halos around its colonies on the same lawn.

**Fig 2.**
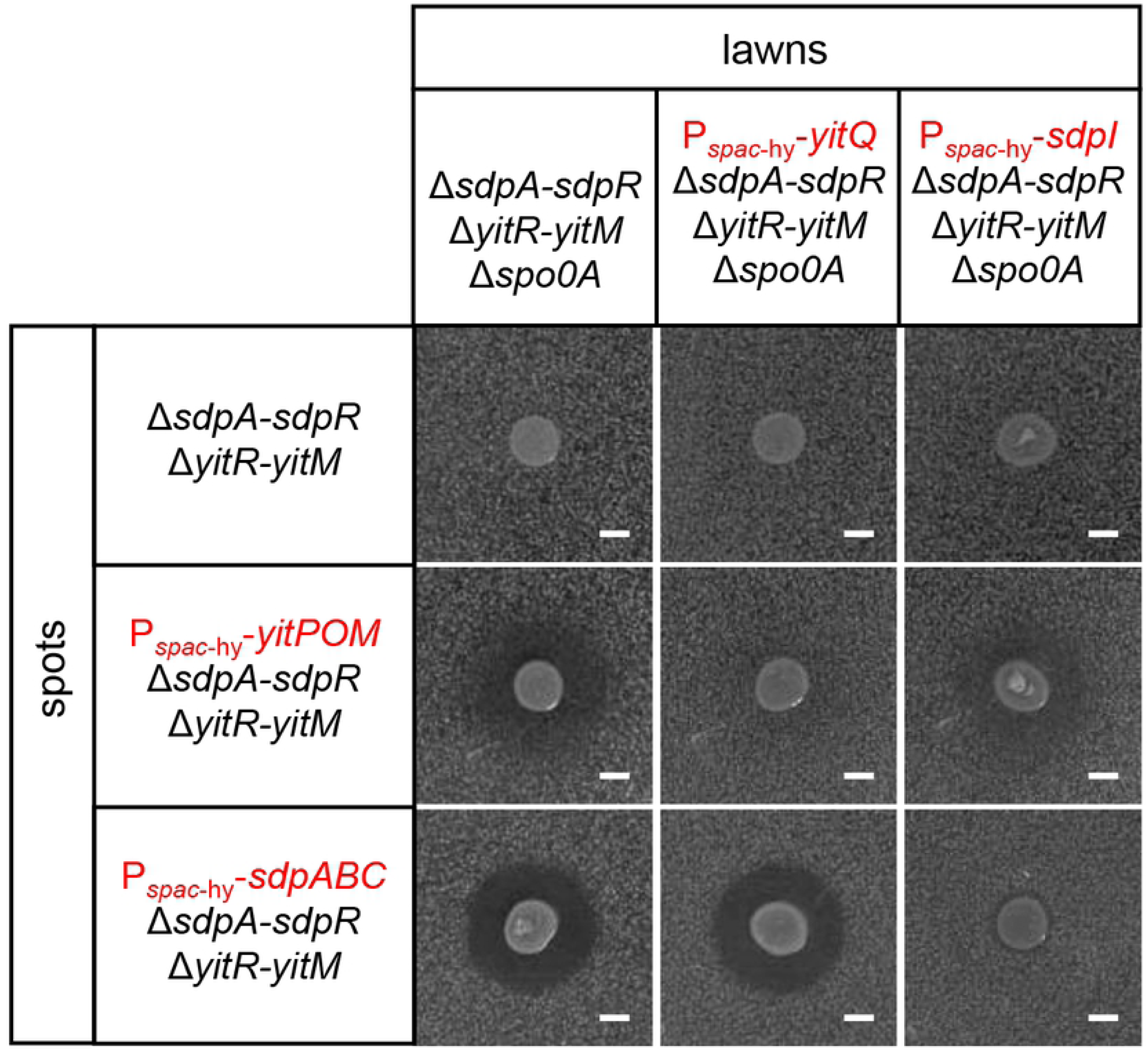
YitQ is an anti-toxin protein to the YIT toxin. *B. subtilis* strains Δ*sdpA-sdpR* Δ*yitR-yitM* Δ*spo0A*, P*_spac_*_-hy_-*yitQ* Δ*sdpA-sdpR* Δ*yitR-yitM* Δ*spo0A*, and P*_spac_*_-hy_-*sdpI* Δ*sdpA-sdpR* Δ*yitR-yitM* Δ*spo0A* were added to 1.2% LB agar and poured as lawns. Strains tested for antibiotic production (shown on the left of the figure as spots) were spotted on the lawns. A growth inhibitory zone was observed if the lawn strain was sensitive to a compound produced by the strain spotted on it. Plates were incubated at 37°C. Scale bar, 2 mm.

These results demonstrate that, like *sdpABC*, *yitPOM* encodes a secreted toxin, which we named YIT.

### YitQ is an anti-toxin protein to the YIT toxin

To determine whether YitQ is an anti-toxin protein to the YIT toxin, we examined the effect of *yitQ* overexpression on the toxin activity of YIT. To this end, the P*_spac_*_-hy_-*yitQ* construct was introduced into the *amyE* locus of the indicator strain (i.e., the Δ*sdpA-sdpR* Δ*yitR-yitM* Δ*spo0A* mutant). When spotted on a lawn of the indicator strain expressing YitQ (P*_spac_*_-hy_-*yitQ* Δ*sdpA-sdpR* Δ*yitR-yitM* Δ*spo0A*), the strain expressing YitPOM (P*_spac_*_-hy_-*yitPOM* Δ*sdpA-sdpR* Δ*yitR-yitM*) did not form halos around its colonies (Fig 2). Thus, *yitQ* expression confers resistance to the YIT toxin.

We were interested in whether there is crosstalk between *yitPOM*/*yitQ* and *sdpABC*/*sdpI*. To explore this possibility, a strain expressing SdpABC (P*_spac_*_-hy_-*sdpABC* Δ*sdpA-sdpR* Δ*yitR-yitM*) was spotted onto lawns of the control indicator strain (Δ*sdpA-sdpR* Δ*yitR-yitM* Δ*spo0A*) and the indicator strain expressing YitQ (P*_spac_*_-hy_-*yitQ* Δ*sdpA-sdpR* Δ*yitR-yitM* Δ*spo0A*) (Fig 2). The SdpABC-expressing strain formed clear halos around its colonies on both types of lawns. Thus, *yitQ* expression did not confer resistance to the SDP toxin. We also tested whether SdpI expression confers resistance to the YIT and SDP toxins. A strain expressing YitPOM (P*_spac_*_-hy_-*yitPOM* Δ*sdpA-sdpR* Δ*yitR-yitM*) formed halos around its colonies on a lawn of the indicator strain expressing SdpI (P*_spac_*_-hy_-*sdpI* Δ*sdpA-sdpR* Δ*yitR-yitM* Δ*spo0A*), whereas the strain expressing SdpABC (P*_spac_*_-hy_-*sdpABC* Δ*sdpA-sdpR* Δ*yitR-yitM*) did not. These results indicate that YitQ and SdpI are anti-toxin proteins specific to the YIT and SDP toxins, respectively. Thus, the two toxin/anti-toxin gene pairs *yitPOM*/*yitQ* and *sdpABC*/*sdpI* most likely function independently.

### Expression of *yitPOM* and *yitQ*

The *yitPOM* operon was previously identified as a member of the DegS-DegU-regulated genes via a DNA microarray analysis using another *B. subtilis* strain, ATCC6051 [48]. To confirm this property in strain 3610, we carried out a Northern blot analysis. RNA samples were isolated from wild-type and Δ*degU* mutant cells grown for various lengths of time in 2×SG with vigorous shaking (Fig 3A). We detected a single band at a position between the 23S rRNA (2904 nt) and 16S rRNA (1541 nt) on Northern blots with a *yitP*-specific probe (Fig 3B). The size of the band was consistent with the length of the entire *yitPOM* locus (2031 bp), confirming that the *yitPOM* locus is transcribed as an operon (Fig 3C). On the Northern blots, the *yitPOM* transcript was observed in the stationary phase samples from the wild-type strain but not in those from the Δ*degU* mutant (Fig 3B). These results indicate that DegS-DegU directly or indirectly induces *yitPOM* transcription in stationary phase.

**Fig 3.**
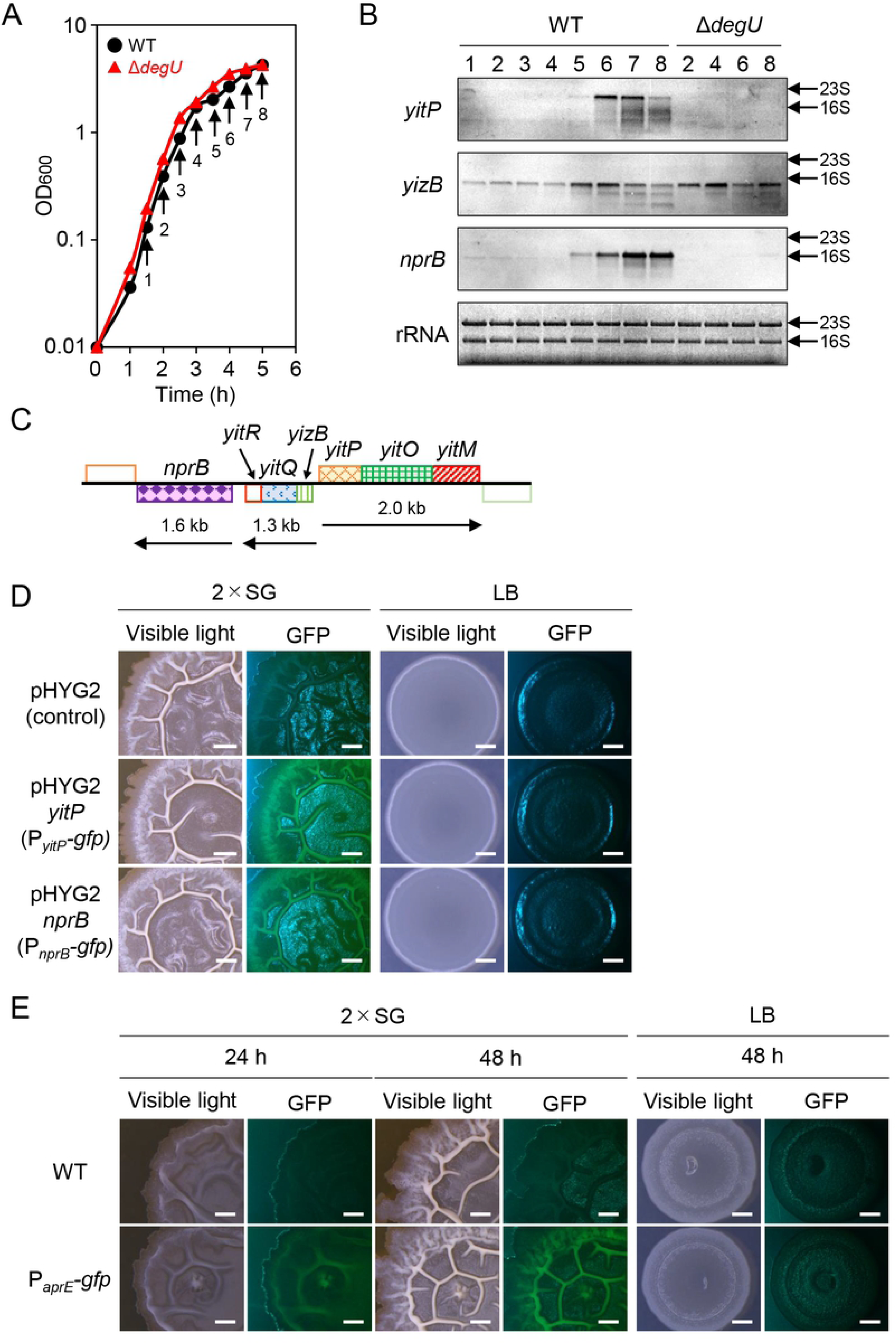
*yitPOM* expression is induced in biofilms by DegS-DegU. (A) Growth profiles of the wild-type and Δ*degU* mutant strains. Strains were grown in 2×SG with vigorous shaking. Arrows indicate the time points at which samples were taken for RNA isolation. (B) Northern blot analysis of *yitPOM*, *yizB-yitQ-yitR*, and *nprB*. Transcripts were detected with gene-specific DIG-labeled RNA probes. Lane numbers (time points) under the strain names correspond to the time points shown in panel A. rRNA stained with methylene blue is shown as a loading control. The positions of 23S rRNA and 16S rRNA are indicated by arrows. (C) Expression of *yitPOM* and *nprB* in biofilms. The wild-type strain carrying the multi-copy plasmid pHYG2 (promoterless *gfp*), pHYG2-*yitP* containing the P*_yitP_*-*gfp* reporter, or pHYG2-*nprB* containing the P*_nprB_-gfp* reporter was grown at 37°C for 24 h on 2×SG or LB. Scale bar, 1 mm. (D) Expression of *aprE* in biofilms. The P*_aprE_*-*gfp* strain was grown at 30°C for 48 h on 2×SG or LB. Colonies of the wild-type strain are shown as a reference. Scale bar, 1 mm.

*yitQ* is predicted to form an operon with its upstream and downstream genes, *yizB* and *yitR*. We detected a band below the position of 16S rRNA on Northern blots with a *yizB*-specific probe (Fig 3B). The size of the band was consistent with the length of the *yizB-yitQ*-*yitR* locus (1244 bp), supporting the conclusion that *yizB*, *yitQ*, and *yitR* are transcribed as an operon (Fig 3C). Based on the Northern blots, the *yizB-yitQ-yitR* operon was transcribed at low levels during exponential phase and then induced during stationary phase in the wild-type strain (Fig 3B). The Δ*degU* mutation had no significant effect on the transcription of the *yizB-yitQ-yitR* operon. The SDP toxin mediates cannibalism between “Spo0A ON” and “Spo0A OFF” cells [40]. The finding that DegS-DegU regulates *yitPOM* but not *yitQ* rules out the possibility that the YIT toxin mediates cannibalism between “DegU ON” and “DegU OFF” cells.

To explore the function of the YIT toxin, we asked under what conditions *yitPOM* expression is induced. The DegS-DegU-regulated gene *bpr*, which encodes an extracellular protease, is expressed in biofilms [56]. We therefore speculated that *yitPOM* is also expressed in biofilms. To visualize *yitPOM* expression in biofilms, the *yitP* promoter was fused to the green fluorescent protein (GFP) reporter, and the resulting P*_yitP_-gfp* reporter construct was introduced to the *amyE* locus on the chromosome of the wild-type strain. *B. subtilis* biofilms are wrinkled structures on the surfaces of colonies grown on solid media that support biofilm formation, such as 2×SG [24]; therefore, we attempted to examine the expression level of the P*_yitP_-gfp* reporter in colonies grown on 2×SG solid medium. We did not detect any fluorescent GFP signal on these colonies, probably because the *yitP* promoter activity was too low. We next examined the expression of the P*_yitP_-gfp* reporter inserted into the multi-copy plasmid pHYG2. The expression of the P*_yitP_-gfp* reporter on the plasmid was examined at 37°C because the plasmid pHYG2 negatively affected biofilm formation at 30°C. The wild-type strain carrying the multi-copy plasmid pHYG2-*yitP* (P*_yitP_-gfp*) formed wrinkled structures on the surfaces of colonies grown on 2×SG solid medium as the biofilms developed. Weak GFP fluorescent signal was observed in these wrinkles (Fig 3D). By contrast, this strain formed flat colonies on LB medium, which does not support biofilm formation, and produced no detectable GFP signal. Under the same conditions, the wild-type strain carrying the parental plasmid pHYG2 (promoterless *gfp*) produced no detectable fluorescent signal on either 2×SG or LB. To eliminate potential artifacts resulting from multi-copy plasmid-based experiments, we examined the expression of *aprE*, which is one of the most highly expressed genes among the DegS-DegU-regulated genes [47]. To this end, we used a strain carrying a single copy of the P*_aprE_-gfp* reporter inserted into the *amyE* locus on the chromosome. The P*_aprE_-gfp* strain produced GFP fluorescent signal in the wrinkles of colonies grown on 2×SG as the wrinkle structures developed (Fig 3E). By contrast, no detectable GFP signal was observed when the P*_aprE_-gfp* strain was grown on LB. These results indicate that DegS-DegU induces its regulatory target genes, including *yitPOM*, in biofilms and that the YIT toxin may play a role in biofilms.

### The YIT toxin inhibits colony biofilm formation

We examined whether *yitPOM* overexpression from the *spac*-hy promoter affects biofilm formation. We used two media, the complex medium 2×SG and the synthetic medium MSgg [24], to support biofilm formation. On 2×SG solid medium, the wild-type strain formed whitish wrinkled colonies (Fig 4A). Induction of *yitPOM* did not affect the colony morphologies of the wild-type, Δ*yitR-yitM* mutant, or Δ*sigW* mutant strains; these P*_spac_*_-hy_-*yitPOM* strains formed similar whitish wrinkled colonies in the presence or absence of IPTG. By contrast, *yitPOM* induction altered the colony morphology of the Δ*yitR-yitM* Δ*sigW* mutant; the P*_spac_*_-hy_-*yitPOM* Δ*yitR-yitM* Δ*sigW* mutant formed brown flat colonies in the presence of IPTG (Fig 4A). Magnified images showed that the whitish wrinkled layers (biofilms) were completely absent on the surfaces of the P*_spac_*_-hy_-*yitPOM* Δ*yitR-yitM* Δ*sigW* mutant colonies (Fig 4B). Similar results were obtained on MSgg medium. The wild-type strain formed light brown wrinkled colonies on MSgg (Fig 4A). Induction of *yitPOM* altered the colony morphology of the Δ*yitR-yitM* Δ*sigW* mutant. The P*_spac_*_-hy_-*yitPOM* Δ*yitR-yitM* Δ*sigW* mutant formed brown flat colonies in the presence of IPTG (Fig 4A), and these colonies completely lacked the light brown wrinkled layers (biofilms) on their surfaces (Fig 4B). Induction of *yitPOM* also altered the colony morphology of the Δ*yitR-yitM* mutant when it was grown on MSgg (Fig 4A). In the presence of IPTG, the P*_spac_*_-hy_-*yitPOM* Δ*yitR-yitM* mutant formed colonies covered with attenuated wrinkles at 96 h post-inoculation; however, these wrinkles faded over time (S2 Fig). We also examined the colony morphologies on the complex medium LB and on the synthetic medium Spizizen minimal medium (SMM) [55], on which *B. subtilis* forms flat colonies rather than biofilms. On these media, *yitPOM* induction had little or no effect on colony morphology, even in the Δ*yitR-yitM* Δ*sigW* mutant (Fig 4A). We compared the effect of *yitPOM* overexpression on colony morphology with that of *sdpABC* overexpression. To this end, *sdpABC* was expressed under the control of the same promoter (*spac*-hy) in the Δ*sdpA-sdpR* Δ*sigW* mutant. The P*_spac_*_-hy_-*sdpABC* Δ*sdpA-sdpR* Δ*sigW* mutant formed normal colonies in all four media in the absence of IPTG, but it did not form colonies in the presence of IPTG (Fig 4A). Thus, the SDP toxin inhibited overall cell growth independently of the medium conditions.

**Fig 4.**
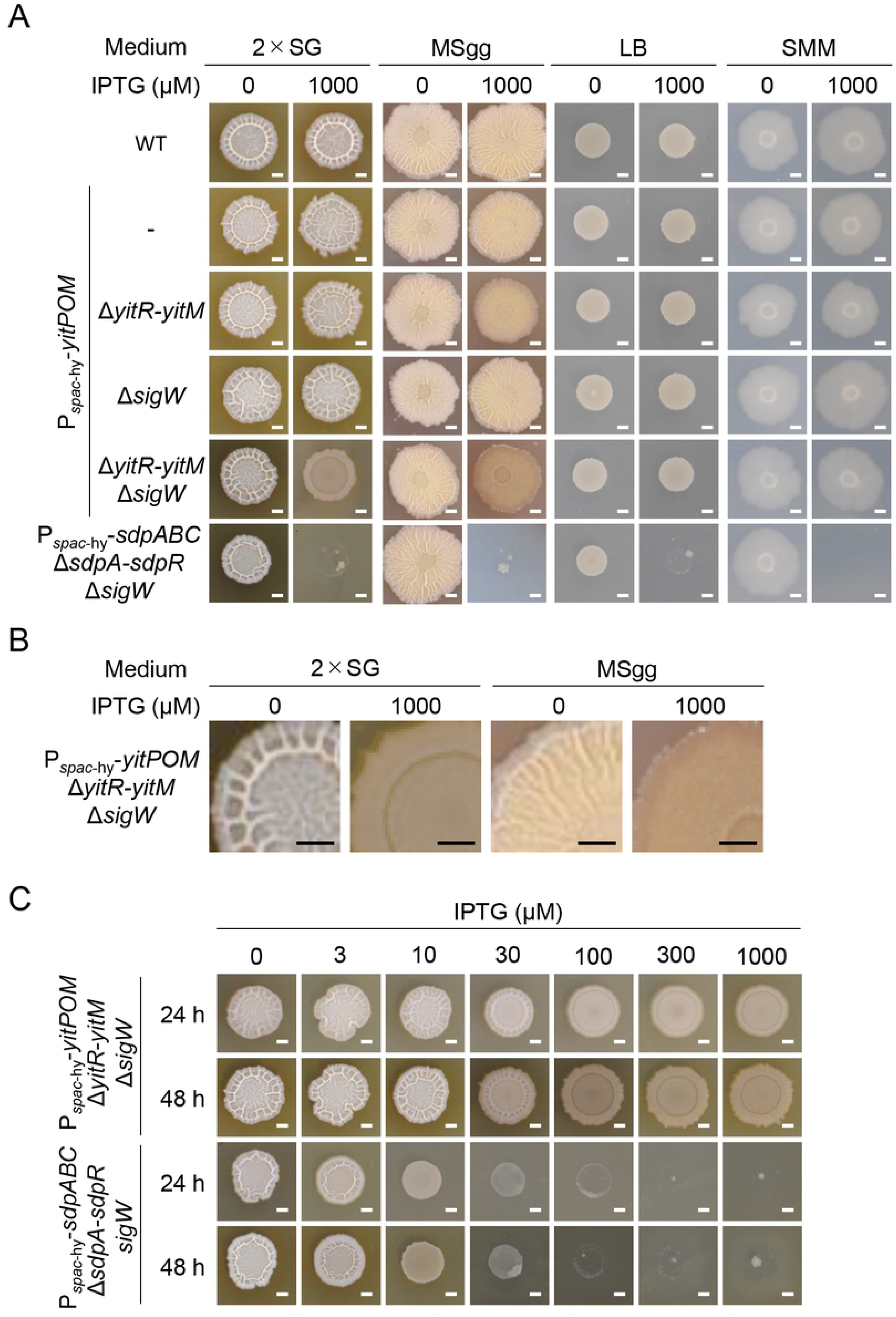
Expression of *yitPOM* inhibits biofilm formation. (A) P*_spac_*_-hy_-*yitPOM* strains with the indicated mutations were grown at 30°C for 48 h on biofilm formation media (2×SG and MSgg) and non-biofilm formation media (LB and SMM) with or without 1 mM IPTG. Colonies of the wild-type and P*_spac_*_-hy_-*sdpABC* Δ*sdpA-sdpR* Δ*sigW* strains are also shown as references. (B) Magnified images of the P*_spac_*_-hy_-*yitPOM* Δ*yitR-yitM* Δ*sigW* mutant colonies shown in panel A. (C) Comparison of the effects of *yitPOM* and *sdpABC* overexpression on colony morphology. The P*_spac_*_-hy_-*yitPOM* Δ*yitR-yitM* Δ*sigW* and P*_spac_*_-hy_-*sdpABC* Δ*sdpA-sdpR* Δ*sigW* mutant strains were grown at 30°C for 48 h on 2×SG with various IPTG concentrations. Scale bar, 2 mm.

We investigated the relationship between the expression levels of *yitPOM* and *sdpABC* and colony morphology. To this end, the P*_spac_*_-hy_-*yitPOM* Δ*yitR-yitM* Δ*sigW* mutant was grown on 2×SG medium supplemented with various IPTG concentrations (0 to 1000 µM) (Fig 4C). The effect of *yitPOM* expression on colony morphology appeared when the P*_spac_*_-hy_-*yitPOM* Δ*yitR-yitM* Δ*sigW* mutant was grown in the presence of 30 µM or higher IPTG concentrations. At 30 µM IPTG, attenuated wrinkles appeared on the colonies at 24 h post-inoculation; however, these wrinkles failed to grow further. At 100 µM IPTG, the colonies completely lacked the whitish wrinkled layers on their surface and became flat. Higher IPTG concentrations did not further alter the colony morphology. Despite having an obvious effect on colony morphology, *yitPOM* induction did not affect colony size. By contrast, when grown on 2×SG medium supplemented with various IPTG concentrations, the P*_spac_*_-hy_-*sdpABC* Δ*sdpA-sdpR* Δ*sigW* mutant formed small colonies in the presence of 10 or 30 µM IPTG but did not form colonies at 100 µM or higher IPTG concentrations (Fig 4C). Thus, *sdpABC* expression exerted a stronger effect on colony formation as its expression levels increased. These results demonstrate that the YIT and SDP toxins have different effects on colony growth and that the YIT toxin specifically inhibits biofilm formation in the absence of its resistance genes.

We also considered how the YIT toxin inhibits biofilm formation. As described above, *yitPOM* induction caused mild cell lysis only in Δ*yitR-yitM* Δ*sigW* mutant cells grown in 2×SG medium with shaking. Induction of *yitPOM* in the Δ*yitR-yitM* Δ*sigW* mutant did not cause cell lysis in cultures grown in LB medium with shaking (S3 Fig). Thus, *yitPOM* induction caused cell lysis and inhibition of biofilm formation only in the Δ*yitR-yitM* Δ*sigW* mutant cells grown on biofilm formation media, indicating that these two phenotypes represent different aspects of one phenomenon. We propose that the YIT toxin likely inhibits biofilm formation by killing biofilm-forming cells rather than by preventing expression of biofilm formation genes.

### NprB allows the YIT toxin to attack cells within biofilms

Induction of *yitPOM* exerted its effects only in cells grown on biofilm formation media. However, the *spac*-hy promoter is active in rich and poor media, including LB and SMM, as observed for the P*_spac_*_-hy_-*sdpABC* Δ*sdpA-sdpR* Δ*sigW* mutant (Fig 4A). These observations suggest the involvement of other factor(s) in the functions of the YIT toxin. A comparison of the genetic organization of the 3610 and BEST195 strains revealed that, in addition to *yizB* and *yitQ*, *nprB* appears to be inserted into the 3610 genome along with *yitPOM* (Fig 1C). *nprB*, which encodes an extracellular neutral protease, was transcribed in a DegU-dependent manner (Fig 3B), and its expression was observed in biofilms (Fig 3D). To determine whether *nprB* is involved in the YIT toxin function, we introduced a deletion of the *nprB-yitM* region (Fig 1C) into the P*_spac_*_-hy_-*yitPOM* Δ*sigW* mutant and examined the colony morphology of the resulting strain. Unlike in the P*_spac_*_-hy_-*yitPOM* Δ*yitR-yitM* Δ*sigW* mutant, *yitPOM* induction did not inhibit biofilm formation in the P*_spac_*_-hy_-*yitPOM* Δ*nprB-yitM* Δ*sigW* mutant (Fig 5A). This strain formed whitish wrinkled colonies like those of the wild-type strain in the presence or absence of IPTG (Fig 5A). Because *nprB* deletion was the only genetic difference between P*_spac_*_-hy_-*yitPOM* Δ*yitR-yitM* Δ*sigW* and P*_spac_*_-hy_-*yitPOM* Δ*nprB-yitM* Δ*sigW* mutants (Fig 1C), this result suggests that the NprB protease is required for the production or function of the YIT toxin.

**Fig 5.**
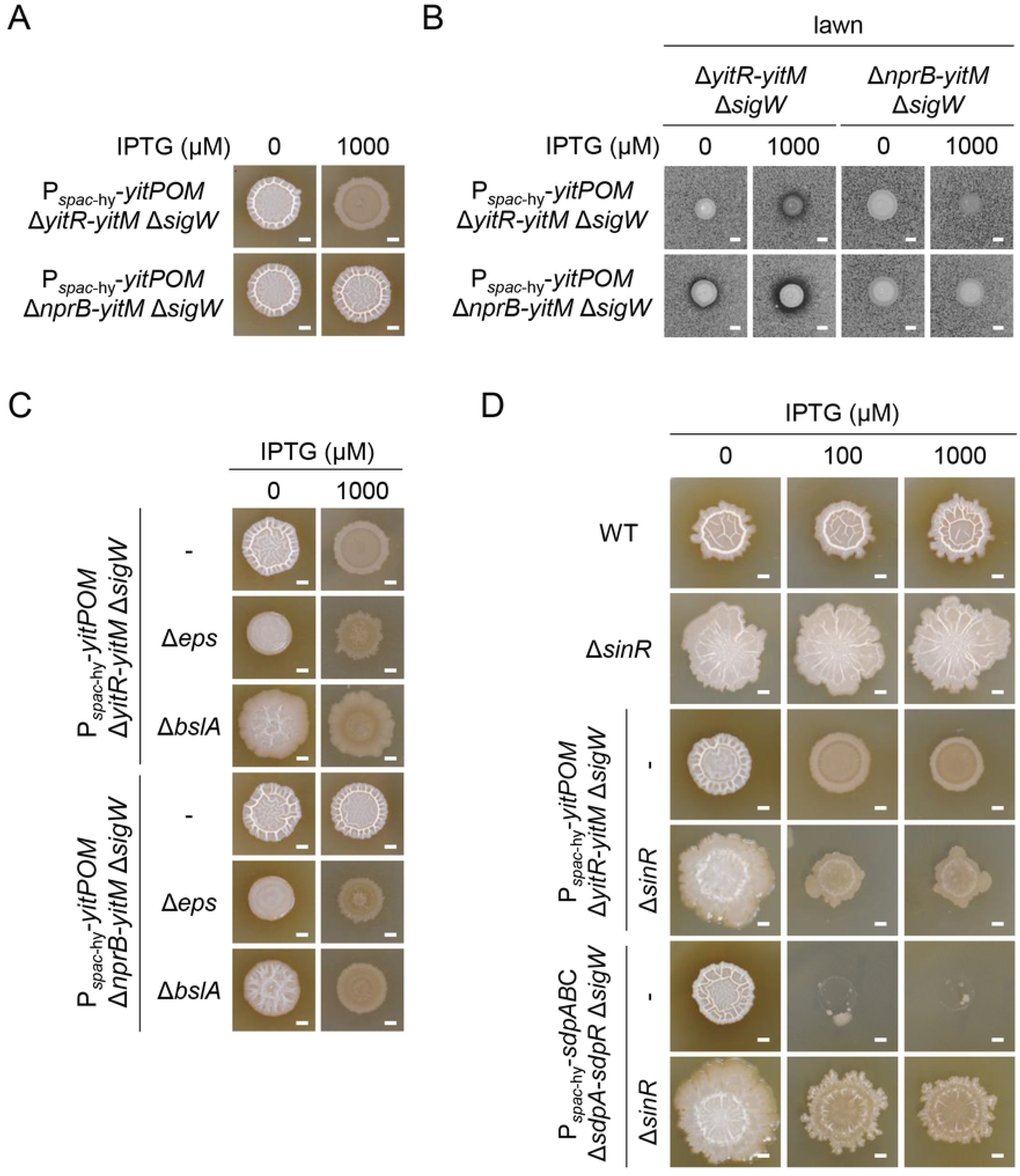
The NprB protease is required for the YIT toxin to inhibit biofilm formation. (A) The Δ*nprB* mutation prevents the YIT toxin from inhibiting biofilm formation. P*_spac_*_-hy_-*yitPOM* Δ*yitR-yitM* Δ*sigW* and P*_spac_*_-hy_-*yitPOM* Δ*nprB-yitM* Δ*sigW* cells were grown on 2×SG at 30°C for 48 h with or without 1000 μM IPTG. (B) Production of the YIT toxin. Δ*yitR-yitM* Δ*sigW* and Δ*nprB-yitM* Δ*sigW* cells were added to 2×SG 1.2% agar with or without 1000 μM IPTG, and the mixtures were poured into plates. P*_spac_*_-hy_-*yitPOM* Δ*yitR-yitM* Δ*sigW* and P*_spac_*_-hy_-*yitPOM* Δ*nprB-yitM* Δ*sigW* cells were spotted on these lawn plates. The plates were then incubated at 37°C for 24 h. (C) The Δ*eps* and Δ*bslA* mutations bypass the requirement for NprB in the ability of the YIT toxin to inhibit biofilm formation. (D) Overproduction of biofilm matrix polymers interfered with the action of the SDP toxin but not with that of the YIT toxin. Scale bar, 2 mm.

To distinguish these possibilities, we examined the production of the YIT toxin in these mutants via spot-on-lawn assays. The P*_spac_*_-hy_-*yitPOM* Δ*yitR-yitM* Δ*sigW* and P*_spac_*_-hy_-*yitPOM* Δ*nprB-yitM* Δ*sigW* mutants were spotted on the lawn of the Δ*yitR-yitM* Δ*sigW* mutant. Although both mutants formed halos around their colonies in the presence of IPTG, the P*_spac_*_-hy_-*yitPOM* Δ*nprB-yitM* Δ*sigW* mutant formed clearer halos than did the P*_spac_*_-hy_-*yitPOM* Δ*yitR-yitM* Δ*sigW* mutant (Fig 5B). The P*_spac_*_-hy_-*yitPOM* Δ*yitR-yitM* Δ*sigW* mutant formed small colonies on the lawn, likely due to loss of biofilm formation. Therefore, we compared the YIT toxin production between the P*_spac_*_-hy_-*yitPOM* Δ*yitR-yitM* and P*_spac_*_-hy_-*yitPOM* Δ*nprB-yitM* mutants. Although these mutants formed similar colonies on the lawn of the Δ*yitR-yitM* Δ*sigW* mutant, the P*_spac_*_-hy_-*yitPOM* Δ*nprB-yitM* mutant formed clearer halos than did the P*_spac_*_-hy_-*yitPOM* Δ*yitR-yitM* mutant (S4 Fig). Thus, the Δ*nprB* mutation increased YIT toxin production. These results suggest that NprB is not required for YIT toxin production or rather that NprB negatively controls the YIT toxin levels within biofilms. The YIT toxin is probably a NprB substrate. We next examined the alternative possibility that the Δ*nprB* mutation might confer resistance to the YIT toxin. To test this idea, we spotted the P*_spac_*_-hy_-*yitPOM* Δ*yitR-yitM* Δ*sigW* and P*_spac_*_-hy_-*yitPOM* Δ*nprB-yitM* Δ*sigW* mutants on the lawn of the Δ*nprB-yitM* Δ*sigW* mutant. Both mutants failed to form clear halos around their colonies (Fig 5B), supporting this idea.

Cells in biofilms are covered with and protected by biofilm matrix polymers, a key reason why cells in biofilms exhibit increased antibiotic tolerance or resistance [5, 6]. We hypothesized that a similar mechanism might work against the YIT toxin and that the NprB protease might enable the YIT toxin molecules to pass through the layers of the biofilm matrix polymers to attack cells within the biofilms. If this were true, then disrupting the biofilm matrix would enable the YIT toxin to inhibit biofilm formation even in the Δ*nprB* mutant. The biofilm matrix of *B. subtilis* biofilms mainly consists of exopolysaccharides (synthesized by the products of the *eps* operon) and polymers of the TasA and BslA proteins [24–29]. To test our hypothesis, we introduced Δ*eps* and Δ*bslA* deletion mutations into the P*_spac_*_-hy_-*yitPOM* Δ*yitR-yitM* Δ*sigW* and P*_spac_*_-hy_-*yitPOM* Δ*nprB-yitM* Δ*sigW* mutants and examined their colony morphologies. The P*_spac_*_-hy_-*yitPOM* Δ*yitR-yitM* Δ*sigW* Δ*eps* mutant formed whitish mucoid colonies in the absence of IPTG, while it formed flat brown colonies in the presence of IPTG (Fig 5C). The difference in colony morphology depending on the presence or absence of IPTG indicates that the induced YIT toxin can function in these colonies even though the Δ*eps* mutation impaired biofilm formation and led to the formation of mucoid colonies. Likewise, the P*_spac_*_-hy_-*yitPOM* Δ*nprB-yitM* Δ*sigW* Δ*eps* mutant also formed whitish mucoid colonies in the absence of IPTG and flat brown colonies in the presence of IPTG (Fig 7B). Thus, the Δ*nprB* mutation did not interfere with the function of the YIT toxin in the Δ*eps* mutant. Similar results were obtained with the Δ*bslA* mutant. The P*_spac_*_-hy_-*yitPOM* Δ*yitR-yitM* Δ*sigW* Δ*bslA* and P*_spac_*_-hy_-*yitPOM* Δ*nprB-yitM* Δ*sigW* Δ*bslA* mutants formed whitish mucoid colonies in the absence of IPTG and flat brown colonies in the presence of IPTG (Fig 5C). These results demonstrate that the Δ*eps* and Δ*bslA* mutations eliminate the requirement for NprB in the function of the YIT toxin. Thus, our idea that NprB enables the YIT toxin to pass through the layers of the biofilm matrix polymers to attack cells in the biofilm is very likely.

To further test this idea, we examined the effect of overexpressing biofilm matrix polymers on the YIT toxin. SinR is a major repressor of the biofilm matrix synthesis genes [32], and a Δ*sinR* mutant formed large swollen colonies due to overproduction of biofilm matrix polymers (Fig 5D). We introduced the Δ*sinR* mutation into the P*_spac_*_-hy_-*yitPOM* Δ*yitR-yitM* Δ*sigW* mutant and examined the colony morphology of the resulting strain. In the absence of IPTG, the P*_spac_*_-hy_-*yitPOM* Δ*yitR-yitM* Δ*sigW* Δ*sinR* mutant formed large swollen colonies, like those of the Δ*sinR* mutant. Induction of *yitPOM* inhibited biofilm formation in this mutant. The P*_spac_*_-hy_-*yitPOM* Δ*yitR-yitM* Δ*sigW* Δ*sinR* mutant formed flat brown colonies in the presence of 100 or 1000 µM IPTG, as was also observed in the P*_spac_*_-hy_-*yitPOM* Δ*yitR-yitM* Δ*sigW* mutant (Fig 5D). We also examined the effect of the Δ*sinR* mutation on the SDP toxin. The P*_spac_*_-hy_-*sdpABC* Δ*sdpA-R* Δ*sigW* Δ*sinR* mutant formed large swollen colonies in the absence of IPTG. Induction of *sdpABC* did not inhibit colony formation and only partly suppressed the swollen colony phenotype even in the presence of 1000 µM IPTG (Fig 5D). These results indicate that overproduction of biofilm matrix polymers interferes with the activity of the SDP toxin but not with that of the YIT toxin. Based on these results, we conclude that the YIT toxin adapts to the biofilm environment and that it can function within mature biofilms with the assistance of the neutral protease NprB.

We examined the colony morphology of the Δ*nprB* mutant. The Δ*nprB* mutant formed wrinkled colonies on 2×SG medium similar to those of the wild-type strain (Fig 6A). We extracted the extracellular proteins and cell surface-associated proteins from these colonies and analyzed them via SDS-PAGE. We detected little or no difference in the protein composition between the wild-type and Δ*nprB* mutant strains in the gels after Coomassie brilliant blue (CBB) staining (Fig 6B). These results indicate that, despite a clear effect on the function of the YIT toxin, the Δ*nprB* mutation does not significantly alter biofilm structure.

**Fig 6.**
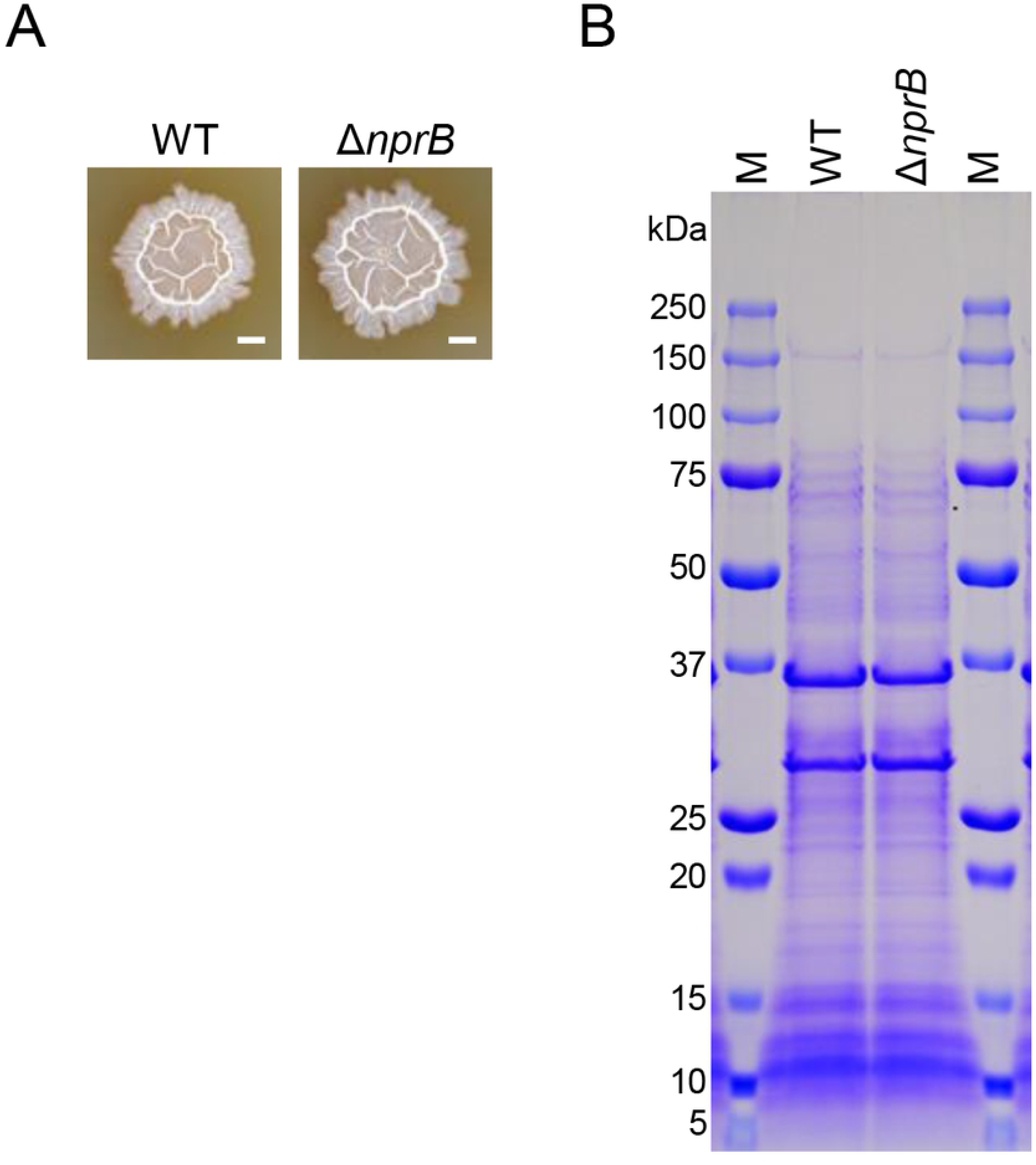
The Δ*nprB* mutation had no significant effect on biofilm formation. (A) Colony biofilms of the wild-type and Δ*nprB* mutant strains. These strains were grown at 30°C for 48 h on 2×SG. Scale bar, 2 mm. (B) The Δ*nprB* mutation had no significant effect on the composition of the extracellular proteins of colony biofilms. Colonies grown at 30°C for 48 h on 2×SG were suspended in SDS-PAGE sample buffer (62.5 mM Tris-HCl (pH 6.8), 1% SDS, 10% glycerol, 2.5% 2-mercaptoethanol, 2 mM PMSF, and 5 mM EDTA) and boiled for 2 min. After centrifugation, the supernatants were subjected to SDS-PAGE. Protein bands were visualized via Coomassie brilliant blue (CBB) staining. The size of protein molecular weight markers (lanes M) is indicated on the left side.

### The YIT toxin functions in the wild-type strain

So far, we have reported the results of experiments designed to uncover the function of the YIT toxin via *yitPOM* expression from the strong *spac*-hy promoter. We asked whether *yitPOM* expression from its own promoter is sufficiently high to exhibit the phenotypes observed above. To address this question, we examined the colony morphologies of mutants lacking *yitQ* and/or *sigW* on 2×SG medium (Fig 7A). While the Δ*yitQ* and Δ*sigW* single mutants formed whitish wrinkled colonies like those of the wild-type strain, the Δ*yitQ* Δ*sigW* double mutant formed colonies with attenuated wrinkles. The Δ*yitR-yitM* Δ*sigW* mutant, which lacks both the toxin and anti-toxin genes, formed whitish wrinkled colonies like those of the wild-type strain. Thus, the phenotype of the Δ*yitQ* Δ*sigW* mutant was caused by the YIT toxin. However, the phenotype of the Δ*yitQ* Δ*sigW* mutant was slightly less noticeable than that of the P*_spac_*_-hy_-*yitPOM* Δ*yitR-yitM* Δ*sigW* mutant in the presence of IPTG.

**Fig 7.**
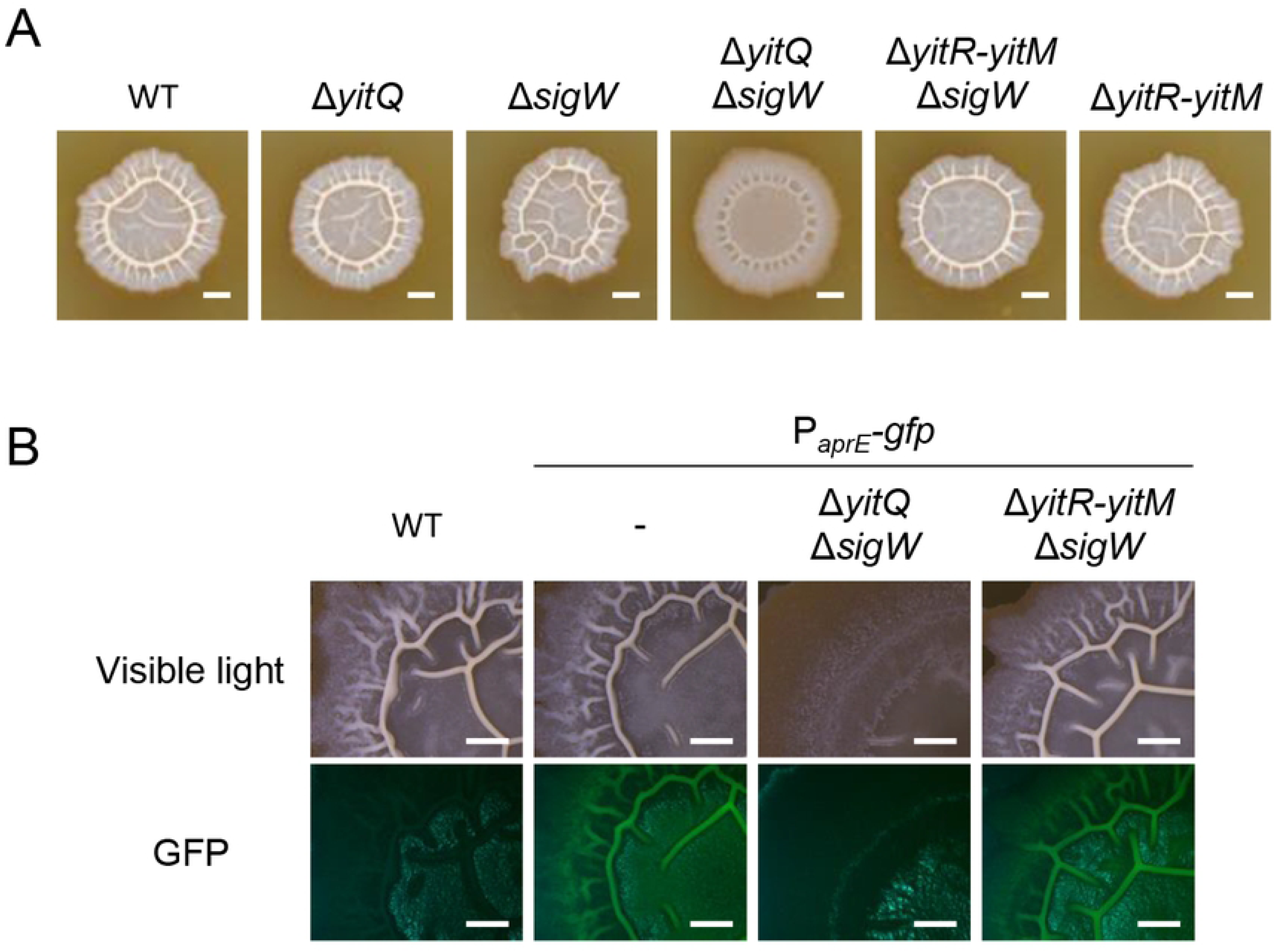
The YIT toxin is expressed and functions in the wild-type strain. (A) Colony morphologies of mutants lacking the resistance genes to the YIT toxin. The strains were grown at 30°C for 48 h on 2×SG. Scale bar, 2 mm. (B) P*_aprE_*-*gfp* expression was reduced in the Δ*yitQ* Δ*sigW* mutant. Scale bar, 1 mm.

As mentioned above, the P*_aprE_-gfp* reporter is expressed in biofilms. The wild-type strain and the Δ*yitR-yitM* Δ*sigW* mutant with the P*_aprE_-gfp* reporter displayed bright GFP signal, while the Δ*yitQ* Δ*sigW* mutant with the P*_aprE_-gfp* reporter displayed no detectable GFP signal on its colonies (Fig 7B). These results indicate that the YIT toxin reduces the number of biofilm-forming cells that express DegS-DegU-regulated genes in the Δ*yitQ* Δ*sigW* mutant. In other words, the action of the YIT toxin reduces the number of cells producing the YIT toxin. This effect could explain why the phenotype of the Δ*yitQ* Δ*sigW* mutant was slightly less obvious. These results demonstrate that the YIT toxin is expressed and functions within biofilms of the wild-type strain.

### The YIT toxin can mediate intercellular competition within biofilms

We asked whether the YIT toxin can mediate intercellular competition within biofilms. To this end, we designed the following experiment. Dilutions of cultures of the wild-type and Δ*yitR-yitM* Δ*sigW* mutant strains were mixed, and the mixtures were spotted on 2×SG solid medium. The inoculated cells grew and formed biofilms in which the wild-type cells were expected to produce the YIT toxin. If the YIT toxin suppressed the growth of Δ*yitR-yitM* Δ*sigW* mutant cells within biofilms, the ratio of these strains within the biofilms would change from the initial ratio. To estimate the ratio of two strains within biofilms, the P*_aprE_-gfp* reporter was introduced into one strain to detect its cells within biofilms.

First, wild-type cells carrying the P*_aprE_-gfp* reporter (the P*_aprE_-gfp* strain) were mixed with wild-type cells at various ratios from 10:0 to 0:10, and the mixtures were spotted on 2×SG solid medium. After 2 days of incubation, we observed the colonies with a fluorescence stereomicroscope. A bright GFP fluorescent signal was detected on colonies grown from the 10:0 mixture, and the GFP signal decreased as the proportion of the P*_aprE_-gfp* strain decreased (Fig 8). When P*_aprE_-gfp*-expressing cells were mixed with Δ*yitR-yitM* Δ*sigW* mutant cells and grown on 2×SG medium, the GFP signal on the colonies also decreased as the proportion of the P*_aprE_-gfp* strain decreased; however, its decrease was moderate compared with that in the former experiment. For example, at a ratio of 3:7, obvious GFP signal was observed on colonies grown from the mixture of the *aprE-gfp* and Δ*yitR-yitM* Δ*sigW* mutant strains but not on colonies grown from the mixture of the *aprE-gfp* and wild-type strains. We also introduced the P*_aprE_-gfp* reporter into the Δ*yitR-yitM* Δ*sigW* mutant. When Δ*yitR-yitM* Δ*sigW* P*_aprE_-gfp* mutant cells were mixed with wild-type cells, no GFP signal was observed, even on the colonies grown from the 9:1 mixture. By contrast, when Δ*yitR-yitM* Δ*sigW* P*_aprE_-gfp* mutant cells were mixed with Δ*yitR-yitM* Δ*sigW* mutant cells, the GFP signal on the colonies decreased as the proportion of the Δ*yitR-yitM* Δ*sigW* P*_aprE_-gfp* mutant cells decreased, as was also the case with the mixture of the *aprE-gfp* and wild-type strains. These results demonstrate that the YIT toxin can mediate intercellular competition within biofilms. We propose that the YIT toxin functions within *B. subtilis* biofilms without being obstructed by the biofilm matrix polymers, thus protecting the biofilms from unfavorable competitors.

**Fig 8.**
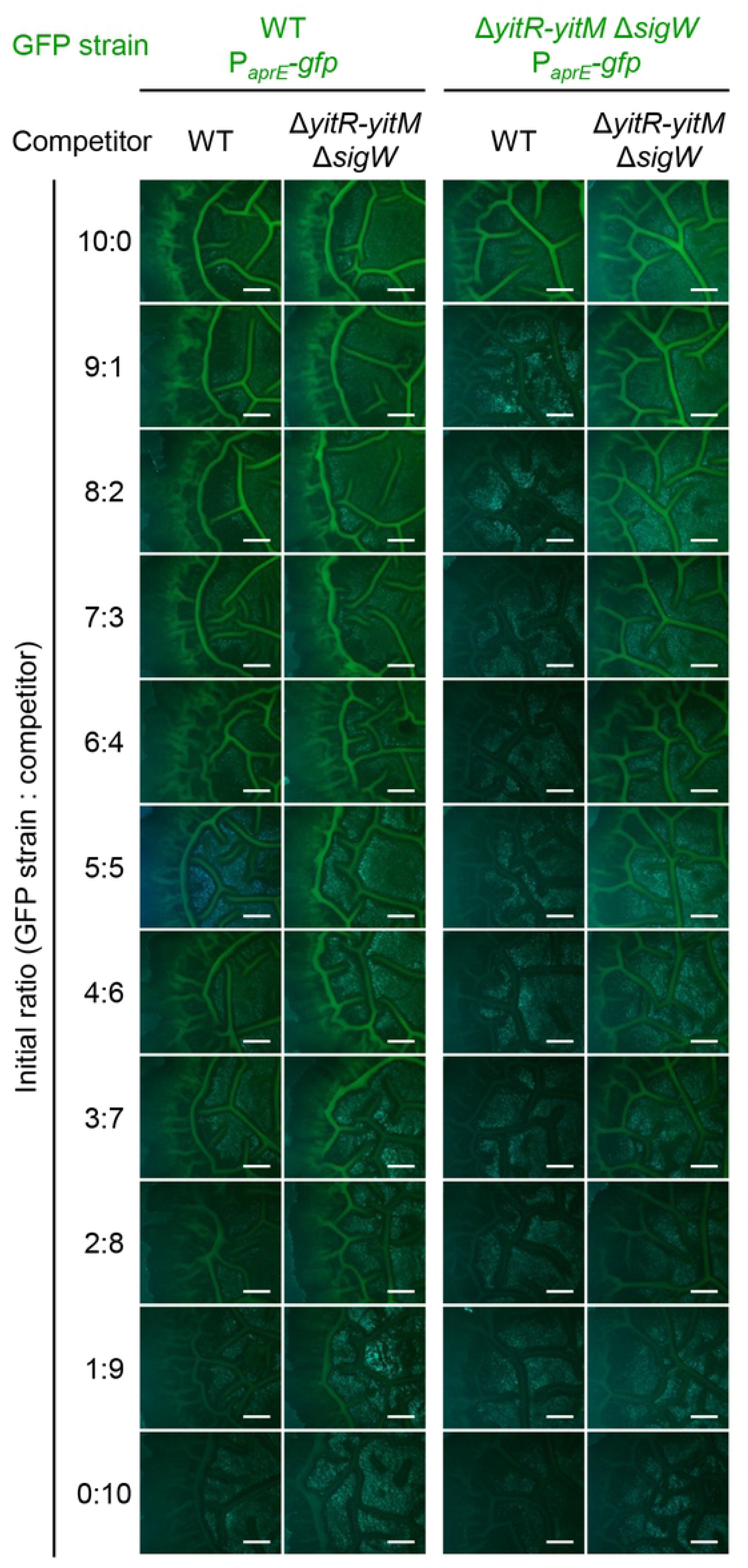
The YIT toxin mediates competition within biofilms. Dilutions of cultures of the indicated strains were mixed at various ratios (10:0 to 0:10) and spotted on 2×SG. P*_aprE_*-*gfp* expression was analyzed 48 h after inoculation. The fluorescent images of colonies are shown. Scale bar, 1 mm.

## Discussion

Biofilms often contain a high-density bacterial community consisting of a mixture of various species, and under these conditions, the bacteria compete with their own siblings and other species for limited space and nutrients. Since cells in biofilms exhibit increased antibiotic tolerance or resistance, the functions of antibiotics in the competition between biofilm bacteria remain unclear. In this study, we demonstrated that *B. subtilis* produces a biofilm-specific toxin, and that this toxin can attack cells within the biofilm by passing through the protective barriers of the biofilm with assistance from an endogenously produced extracellular protease.

The *yitPOM* operon is a paralog of the *sdpABC* operon; however, these operons are expressed in different growth phases. Transcription of *sdpABC* is induced by low levels of Spo0A-P [44]. Given that low Spo0A-P levels also induce the expression of biofilm matrix synthesis genes [44, 45], it is likely that the SDP toxin is induced during the early phase of biofilm formation. By contrast, *yitPOM*, which we showed is activated in biofilms, is induced by DegS-DegU. Moreover, induction of these operons had different effects on *B. subtilis* cells lacking genes whose products confer resistance to the SDP and YIT toxins, i.e., *sdpABC* induction prevented colony formation independently of the medium conditions, whereas *yitPOM* specifically inhibited biofilm formation. Like SdpC, YitM contains a C-terminal hydrophobic domain that seems to be processed to produce the YIT toxin. However, the sequence of the hydrophobic domain of YitM is different from that of SdpC. We assume that this difference contributes to the differences in the functions of these toxins. To summarize, although the *sdpABC* and *yitPOM* operons are paralogous, they have distinct regulations and roles. Indeed, the genomes of many *B. subtilis* strains have both the *sdpABC* and *yitPOM* operons (S1 Table), suggesting that having both *sdpABC* and *yitPOM* may provide survival advantages in the environment. Interestingly, some *B. subtilis* strains possess *sdpABC* homologs that appear to be different from *sdpABC* or *yitPOM* (S1 Table). *sdpABC* homologs, including *sdpABC* itself and *yitPOM*, can be classified into five groups based on their genome position and SdpC homolog sequences (S1 Table, S5 and S6 Figs). Each group of SdpC homologs has a unique C-terminal hydrophobic domain that differs from those of the others (S6 Fig); therefore, various SdpC homolog-derived toxins may have different sequences. If the differences in these sequences impart their functional differences, as observed for SdpC and YitM, then SdpC homolog-derived toxins are likely to play more diverse roles.

Overproduction of biofilm matrix polymers interfered with the function of the SDP toxin but not with that of the YIT toxin, suggesting that the YIT toxin has a mechanism to pass through the layers of the biofilm matrix polymers, and we showed that this mechanism involves the extracellular protease NprB. The YIT toxin could not inhibit biofilm formation in the absence of NprB even though the Δ*nprB* mutation increased the production of the YIT toxin. The requirement of NprB was eliminated by the Δ*eps* and Δ*bslA* mutations, both of which impair biofilm matrix formation and, thus, biofilm formation [24, 28]. Cells in biofilms are encased in the biofilm matrix, which functions as a physical and chemical barrier against antibiotics by limiting their penetration [11–16]. Our results suggest that similar mechanisms might contribute to resistance to the YIT toxin in *B. subtilis* biofilms and that the extracellular protease NprB is required for the YIT toxin to pass through these defense barriers. The Δ*nprB* mutation had no significant effect on the composition of the extracellular and cell surface-associated proteins of biofilms nor on biofilm formation. NprB probably degrades specific proteins in the biofilm matrix to allow the YIT toxin to pass through the layers of the biofilm matrix polymers to attack other biofilm cells.

Although NprB-mediated structural changes in biofilms probably increase the risk that biofilms become susceptible to antibiotics produced by other bacteria, the capability of the YIT toxin to attack sensitive cells within biofilms must be important for maintaining biofilm communities. Biofilms often consist of multiple types and multiple species of bacterial cells, some of which exploit others as free-loaders or cheaters that do not produce biofilm matrix polymers and other public goods [57]. The production of biofilm matrix polymers is metabolically costly; however, biofilm matrix polymers are extracellular products that are accessible even to non-producing cheater cells from which they receive protection [57]. An increase in the number of cheater cells, therefore, can disturb the cooperative relationships within biofilm communities and lead to instability within biofilm communities. The production of antibiotics that can diffuse through the biofilm offers the great advantage of being able to eliminate cheater cells and other unfavorable competitors present in the biofilm. Further work is required to determine which types of cells or what bacterial species are susceptible to the YIT toxin.

Bacterial competition is mediated by multiple factors [58]. In addition to the YIT toxin, DegS-DegU directly or indirectly induces non-ribosomally synthesized peptide antibiotics, e.g., bacilysin, fengycin, iturin, difficidin, and bacillomycin, although the repertoire of antibiotic synthesis genes differs from strain to strain and no *B. subtilis* strain produces all of these [59, 60, 61, 62]. DegS-DegU also induces a wide array of extracellular degradative enzymes, including six extracellular proteases [47 and references therein]. Although these proteases have been thought to play roles in nutrient acquisition from the surrounding environment, our results suggest that these degradative enzymes may also play roles in competition within biofilms. Indeed, several proteases were previously shown to disrupt biofilms of heterologous bacteria [63, 64, 65, 66]. Simultaneously producing multiple antibiotics and degradative enzymes within biofilms affords *B. subtilis* the ability to attack competitor cells protected by their own biofilm matrixes and to exclude them from biofilm communities. We expect that antibiotics and degradative enzymes cooperate extensively in *B. subtilis* biofilms.

Previous studies showed that positively charged antibiotics interact with negatively charged matrix components, such as extracellular DNA and exopolysaccharides, and impede their penetration into biofilms [12, 15]. The SDP toxin contains two positively charged amino acid residues, whereas the hydrophobic region of YitM contains four negatively charged amino acid residues but no positively charged amino acid residues. These observations suggest that both NprB function and the amino acid sequence of the YIT toxin may be important for the ability of the YIT toxin to pass through the layers of biofilm matrix polymers; however, we have not yet determined the relevant sequence of the YIT toxin. The YIT toxin/NprB system seems to have evolved to be specifically adapted to the *B. subtilis* biofilm environment. Some bacteria produce specific antibiotics in biofilms [17–21]. Among them, the *Escherichia coli* ROAR029 strain produces the bacteriocin colicin R in biofilms, and colicin R is more active against biofilms than against planktonic cultures, as is the YIT toxin [20]. *Pseudomonas aeruginosa* produces bacteriocins pyocins and can suppress the growth of pyocin-sensitive bacteria in biofilms [21, 22]. Based on our results and these previous observations, we propose that bacteria have evolved specialized antibiotics that function in biofilms as biofilm-specific competition mechanisms. The properties of these antibiotics may differ from those of conventional antibiotics. Biofilm-associated antibiotics might serve as anti-biofilm agents, especially in combination with degradative enzymes.

## Materials and methods

### Bacterial strains and culture condition

*B. subtilis* strain NCIB3610 and its derivatives used in this study are listed in Table 1. Construction of the *B. subtilis* mutants is described in S1 Appendix. Primers used for the strain construction are listed in S2 Table. *B. subtilis* strains were maintained in LB (LB Lennox; BD Difco, Franklin Lakes, NJ, USA). For colony morphology observation, a fresh single colony was suspended in 100 µl of LB, and 2 µl of the suspension was spotted onto 2×SG [49], MSgg [24], LB, and SMM media [55]. The plates were incubated at 30°C. Colony morphology was observed after 48 h of incubation on 2×SG and LB, after 72 h on SMM, and after 144 h on MSgg.

**Table 1.** *B. subtilis* strains used in this study.

| Strain | Relevant feature(s) | References or construction<sup>a</sup> |
| --- | --- | --- |
| NCIB3610 | prototroph | 24 |
| N1285 | *amyE::P<sub>spac-hy</sub>-yitPOM (erm)* | This study |
| N1286 | *amyE::P<sub>spac-hy</sub>-yitPOM (erm) ΔyitR-yitM::tet* | This study |
| NTF88 | *ΔsigW::cat* | 68 |
| N1356 | *amyE::P<sub>spac-hy</sub>-yitPOM (erm) ΔsigW::cat* | This study |
| N1357 | *amyE::P<sub>spac-hy</sub>-yitPOM (erm) ΔyitR-yitM::tet ΔsigW::cat* | This study |
| N1263 | *ΔyitR-yitM::tet* | This study |
| N1333 | *amyE::P<sub>spac-hy</sub>-sdpABC (erm)* | This study |
| N1335 | *amyE::P<sub>spac-hy</sub>-sdpABC (erm) ΔsdpA-sdpR::spc* | This study |
| N1337 | *amyE::P<sub>spac-hy</sub>-sdpABC (erm) ΔsigW::cat* | This study |
| N1340 | *amyE::P<sub>spac-hy</sub>-sdpABC (erm) ΔsdpA-sdpR::spc ΔsigW::cat* | This study |
| N1458 | *ΔsdpA-sdpR::spc* | This study |
| N741 | *ΔsdpA-sdpR::spc ΔyitR-M::tet* | This study |
| N771 | *ΔsdpA-sdpR::spc ΔyitR-M::tet Δspo0A::cat* | NTF97 [68] → N741 |
| N1459 | *amyE::P<sub>spac-hy</sub>-yitQ (erm) ΔsdpA-sdpR::spc ΔyitR-M::tet Δspo0A::cat* | This study |
| N1460 | *amyE::P<sub>spac-hy</sub>-sdpI (erm) ΔsdpA-sdpR::spc ΔyitR-M::tet Δspo0A::cat* | This study |
| N942 | *amyE::P<sub>spac-hy</sub>-yitPOM (erm) ΔsdpA-sdpR::spc ΔyitR-yitM::tet* | This study |
| N776 | *amyE::P<sub>spac-hy</sub>-sdpABC (erm) ΔsdpA-sdpR::spc ΔyitR-yitM::tet* | This study |
| NTF28 | *ΔdegU::cat* | 62 |
| N1443 | pHYG2 (promoter-less *gfp*, *tet*) | This study |
| N1444 | pHYG2-*yitP* (*P<sub>yitP</sub>-gfp*, *tet*) | This study |
| N1446 | pHYG2-*nprB* (*P<sub>nprB</sub>-gfp*, *tet*) | This study |
| N1382 | *amyE::P<sub>aprE</sub>-gfp (cat)* | W740 [68] → NCIB3610 |
| N1230 | *amyE::P<sub>spac-hy</sub>-yitPOM (erm) ΔyitR-yitM::tet ΔsigW::neo* | This study |
| N1253 | *amyE::P<sub>spac-hy</sub>-yitPOM (erm) ΔyitR-yitM::tet ΔsigW::neo ΔepsA::spc* | This study |
| N1255 | *amyE::P<sub>spac-hy</sub>-yitPOM (erm) ΔyitR-yitM::tet ΔbslA::spc* | This study |
| N1238 | *amyE::P<sub>spac-hy</sub>-yitPOM (erm) ΔnprB-yitM::tet ΔsigW::neo* | This study |
| N1293 | *amyE::P<sub>spac-hy</sub>-yitPOM (erm) ΔnprB-yitM::tet ΔsigW::neo Δeps::spc* | This study |
| N1294 | *amyE::P<sub>spac-hy</sub>-yitPOM (erm) ΔnprB-yitM::tet ΔbslA::spc* | This study |
| N1358 | *amyE::P<sub>spac-hy</sub>-yitPOM (erm) ΔyitR-yitM::tet ΔsigW::neo ΔsinR::cat* | This study |
| N999 | *amyE::P<sub>spac-hy</sub>-sdpABC (erm) ΔsdpA-sdpR::spc ΔsigW::neo* | This study |
| N1334 | *amyE::P<sub>spac-hy</sub>-sdpABC (erm) ΔsdpA-sdpR::spc ΔsigW::neo ΔsinR::cat* | This study |
| N1264 | *nprB::cat* | W115 [47] → NCIB3610 |
| N1287 | *ΔyitQ::cat* | This study |
| N1268 | *ΔsigW::neo* | This study |
| N1288 | *ΔyitQ::cat ΔsigW::neo* | This study |
| N1234 | *ΔyitR-yitM::tet ΔsigW::neo* | This study |
| N1413 | *ΔyitQ::spc ΔsigW::neo amyE::P<sub>aprE</sub>-gfp (cat)* | This study |
| N1388 | *ΔyitR-yitM::tet ΔsigW::neo amyE::P<sub>aprE</sub>-gfp (cat)* | This study |

<sup>a</sup>Arrows indicate *B. subtilis* transformation: donor strain → recipient strain.

### Northern blot analysis

Wild-type and Δ*degU* mutant cells were grown at 37°C in 2×SG with vigorous shaking, and samples were taken from the cultures at various time points for RNA isolation. Total RNA was prepared as previously described [47]. The Northern blot analysis was carried out as previously described [47]. Primers used for RNA probe synthesis are shown in S2 Table.

### Comparison of genetic organization

A comparison of the genetic organization in different *B. subtilis* strains was carried out using the MBGD website (http://mbgd.genome.ad.jp/) [67]. Protein alignments were carried out using the Protein BLAST program on the NCBI website (https://www.nlm.nih.gov/).

### Spot-on-lawn assay

Indicator strains (lawn strains) were grown at 28°C overnight in LB with vigorous shaking. Culture (1 µl) was mixed with 12 ml of 50°C LB 1.2% agar or 2×SG 1.2% agar with brief vortexing, and the mixture was immediately poured into a ϕ9 cm plate. The lawn plates were dried for 20 min in a laminar flow cabinet. The strains tested for antibiotic production were grown at 28°C overnight in LB with vigorous shaking. These cultures were diluted 10 times with LB, and 2 µl of the dilutions were spotted onto the dried lawn plates. The plates were incubated at 37°C for 24 to 30 h until halos appeared around the colonies.

### Microscopic observation

The strains carrying the multicopy *gfp* reporters were grown at 37°C on 2×SG or LB solid medium. The P*_aprE_-gfp* reporter strains were grown at 30°C on 2×SG or LB solid medium. The expression of the GFP reporters on the colonies was analyzed with a SZX7 stereomicroscope (Olympus, Tokyo, Japan) equipped with an AdvanCam-E3Rs digital color camera (Advan Vision, Tokyo, Japan). Images were obtained and processed with AdvanView (Advan Vision) and Photoshop Elements (Adobe, San Jose, CA, USA).

### Competition assay

Strains were grown at 28°C overnight in LB with vigorous shaking. Cultures of the strains used for the assay contained 3.0 × 10^8^ cells/ml on average. The cultures were diluted 100-fold in LB, and two dilutions were mixed at the indicated ratios. Aliquots (2 µl each) of the mixtures were spotted onto 2×SG solid medium, and the plates were incubated at 30°C for 48 h. GFP fluorescence on the colonies was analyzed as described above.

## ACKNOWLEDGMENTS

We would like to thank Prof. Hisaji Maki and Associate Prof. Masahiro Akiyama for their helpful advice and support.

## Supporting Information Legends

**S1 Fig.**

***yitPOM* is a paralog of *sdpABC.***

(A) Similarity between SdpABC and YitPOM. (B) Alignment of SdpC and YitM. The signal sequences and the SDP toxin sequence are shown in blue and red, respectively. Identical and similar amino acids shared by the two proteins are indicated by asterisks and dots, respectively. (C) Hydropathy plots of SdpC and YitM. The plots were constructed using the ExPASy website (https://web.expasy.org/protscale/) with a window size of 19.

**S2 Fig.**

**The colony morphology of P*_spac-_*_hy_-*yitPOM* mutants on MSgg medium with 1 mM IPTG.**

The strains were grown at 30°C on MSgg. Scale bar, 2 mm.

**S3 Fig.**

**The growth profile of the P*_spac_*_-hy_-*yitPOM* Δ*yitR-yitM* Δ*sigW* mutant in LB with or without 1 mM IPTG.**

**S4 Fig.**

**The production of the YIT toxin.**

Δ*yitR-yitM* Δ*sigW* and Δ*nprB*-*yitM* Δ*sigW* cells were added to 1.2% 2×SG agar with or without 1000 μM IPTG, and the mixtures were poured into plates. P*_spac_*_-hy_-*yitPOM* Δ*yitR-yitM* and P*_spac_*_-hy_-*yitPOM* Δ*nprB-yitM* cells were spotted on the lawn plates. The plates were incubated at 37°C for 24 h.

**S5 Fig.**

**Comparison of the genetic organization of the *sdpABC* homologs in different *B. subtilis* strains.**

The genetic organization of three *sdpABC* homologs in the indicated *B. subtilis* strains was compared with that of the corresponding locus in strain NCIB3610. Homologous genes are shown by boxes of the same color. Strain names are shown to the right of the gene maps.

**S6 Fig.**

**Comparison of SdpC homologs.**

(A) The alignment of SdpC homologs. The sequences of SdpC homolog 1 (YitM), homolog 2, homolog 3, and homolog 4 are derived from *B. subtilis* strains NCIB3610, ATCC13952, BEST195, and OH131.1, respectively. The signal sequences and the SDP toxin sequence are shown in blue and red, respectively. Identical and similar amino acids among all of the homologs are indicated by asterisks and dots, respectively. (B) Hydropathy plots of SdpC homologs. The plots were constructed using the ExPASy website (https://web.expasy.org/protscale/) with a window size of 19.

